# Short-term plasticity of the human adult visual cortex measured with 7T BOLD

**DOI:** 10.1101/323741

**Authors:** Paola Binda, Jan W. Kurzawski, Claudia Lunghi, Laura Biagi, Michela Tosetti, Maria Concetta Morrone

**Author notes:** co-first authors (Paola Binda, Jan W. Kurzawski). Lead contact and corresponding author: Maria Concetta Morrone, *University of Pisa, Department of Translational Research and New Technologies in Medicine and Surgery*, Via San Zeno 31, 56123 Pisa (PI), Italy.

## Abstract

Visual cortex, particularly V1, is considered to be resilient to plastic changes in adults. In particular, ocular dominance is assumed to be hard-wired after the end of the critical period. We show that short-term (2h) monocular deprivation in adult humans boosts the BOLD response to the deprived eye, changing ocular dominance of V1 vertices, consistently with homeostatic plasticity. The boost is strongest in V1, present in V2, V3 & V4 but absent in V3a and MT. Assessment of spatial frequency tuning in V1 by a population Receptive-Field technique shows that deprivation primarily boosts high spatial frequencies, consistent with a primary involvement of the parvocellular pathway. Crucially, the V1 deprivation effect correlates across participants with the perceptual increase of the deprived eye dominance assessed with binocular rivalry, suggesting a common origin. Our results demonstrate that visual cortex, particularly the ventral pathway, retains a high potential for homeostatic plasticity in the human adult.

## Introduction

To interact efficiently with the world, our brain needs to fine-tune its structure and function, adapting to a continuously changing external environment. This key property of the brain, called *neuroplasticity*, is maximal early in life, within the so called *critical period* (Berardi, Pizzorusso, & Maffei, 2000; Hubel & Wiesel, 1970; Hubel, Wiesel, & LeVay, 1977; Wiesel & Hubel, 1963), but it is thought to decline dramatically during adulthood, especially at the level of the primary sensory cortices. During development, the plastic cortical response to abnormal visual experience is so strong that occluding one eye for a few days induces a dramatic and permanent shift in ocular dominance (the amount of V1 neurons responding to each eye) in favor of the open eye (Berardi et al., 2000; Gordon & Stryker, 1996; Hubel & Wiesel, 1970; Hubel et al., 1977; Wiesel & Hubel, 1963), while the deprived eye becomes functionally blind or very weak, a phenomenon known as amblyopia (Gordon & Stryker, 1996; Kiorpes, Kiper, O’Keefe, Cavanaugh, & Movshon, 1998; Levi & Carkeet, 1993; Wiesel & Hubel, 1963). After the closure of the critical period, V1 is thought to become fundamentally hard-wired (Mitchell & Sengpiel, 2009; Sato & Stryker, 2008) – although recent evidence in both cat and mice indicates that plasticity can be restored in adults by manipulating visual cortex excitation (Fong, Mitchell, Duffy, & Bear, 2016; Fregnac, Shulz, Thorpe, & Bienenstock, 1988; Maya Vetencourt et al., 2008).

In adult humans, like in the animal model, cortical processes appear to be characterized more by stability than plasticity (Baseler et al., 2002; Baseler et al., 2011; Wandell & Smirnakis, 2009). However, there are at least two phenomena which consistently evoke perceptual plasticity in adults: perceptual learning (B. Dosher & Lu, 2017; Fahle & Poggio, 2002; Fiorentini & Berardi, 1980; A Karni & Sagi, 1991; A. Karni & Sagi, 1993; Watanabe & Sasaki, 2015) and short-term visual deprivation (Binda & Lunghi, 2017; Lunghi, Berchicci, Morrone, & Di Russo, 2015; Lunghi, Burr, & Morrone, 2011; Lunghi, Burr, & Morrone, 2013; Zhang, Bao, Kwon, He, & Engel, 2009 Lunghi, 2015 #35; Zhou, Clavagnier, & Hess, 2013; Zhou, Reynaud, & Hess, 2014). Short-term deprivation in adults typically induces a paradoxical boost of the perception of the stimulus from the deprived eye. A few hours deprivation of one cardinal orientation enhances sensitivity to the deprived orientation (Zhang et al., 2009); similarly, two hours of monocular contrast deprivation leads to a transient boost of the deprived eye (Binda & Lunghi, 2017; Lunghi, Berchicci, et al., 2015; Lunghi et al., 2011; Lunghi et al., 2013; Lunghi, Emir, Morrone, & Bridge, 2015; Zhou et al., 2013; Zhou et al., 2014). The effect of short-term monocular deprivation is long-lasting, particularly for chromatic equiluminant stimuli optimized for stimulation of the parvocellular pathway, for which a significant boost is still observed up to 3h after the end of the short-term deprivation (Lunghi et al., 2013). In addition, the effect of 2h of deprivation can be retained across 6h of sleep (Menicucci, Lunghi, Zaccaro, Morrone, & Gemignani, 2018) and can lead to permanent visual changes in amblyopic patients (Lunghi et al., 2016). All this evidence strongly suggests the involvement of a plastic reorganization of visual cortical processes.

The boost of the deprived eye after short-term monocular deprivation is consistent with homeostatic plasticity, an initial compensatory reaction of the visual system to deprivation aimed at maintaining the average cortical activity constant despite the impoverished incoming visual input (G. Turrigiano, 2012). Homeostatic plasticity was first reported in animal models during the developmental critical period, following many days of monocular deprivation (Mrsic-Flogel et al., 2007; Ranson, Cheetham, Fox, & Sengpiel, 2012; G. Turrigiano, 2012; G. G. Turrigiano & Nelson, 2004). Recently, a form of homeostatic plasticity after short-term monocular deprivation has been observed in adult monkey V1: ocular dominance columns change expanding the representation of the deprived eye (Begum & Tso, 2016; Tso, Miller, & Begum, 2017). Interestingly, like for adult human perception, the macaque V1 deprivation effect is strongest when deprivation mainly affects the parvocellular activity (Begum & Tso, 2016).

In adult humans, the neural substrates of short-term monocular deprivation effects have been indirectly studied with MR spectroscopy (showing a GABA concentration change in the occipital cortex, Lunghi, Emir, et al., 2015) and Visual Evoked Potentials (showing a modulation of the early visual response components, Lunghi, Berchicci, et al., 2015). Here we directly measure the changes in early visual cortical areas using 7T fMRI in adult humans, before and after two hours of monocular deprivation. Assessing the BOLD change and its selectivity to spatial frequency with a newly developed approach (conceptually similar to the population Receptive Field method, Dumoulin & Wandell, 2008), we demonstrate a change of ocular drive of BOLD signals in primary visual cortex, selective for the higher spatial frequencies and strongest along the ventral pathway, consistent with a stronger plasticity potential of the parvocellular pathway in adulthood.

## Results

### Monocular deprivation boosts V1 responses to the deprived eye and shifts BOLD ocular dominance

To investigate the visual modulation of BOLD signal by short term deprivation, we performed ultra-high field (UHF, 7T) fMRI during the presentation of high contrast dynamic visual stimuli, delivered separately to the two eyes, before and after 2h of monocular contrast deprivation (see schematic diagram in Fig. 1A).

**Figure 1:**
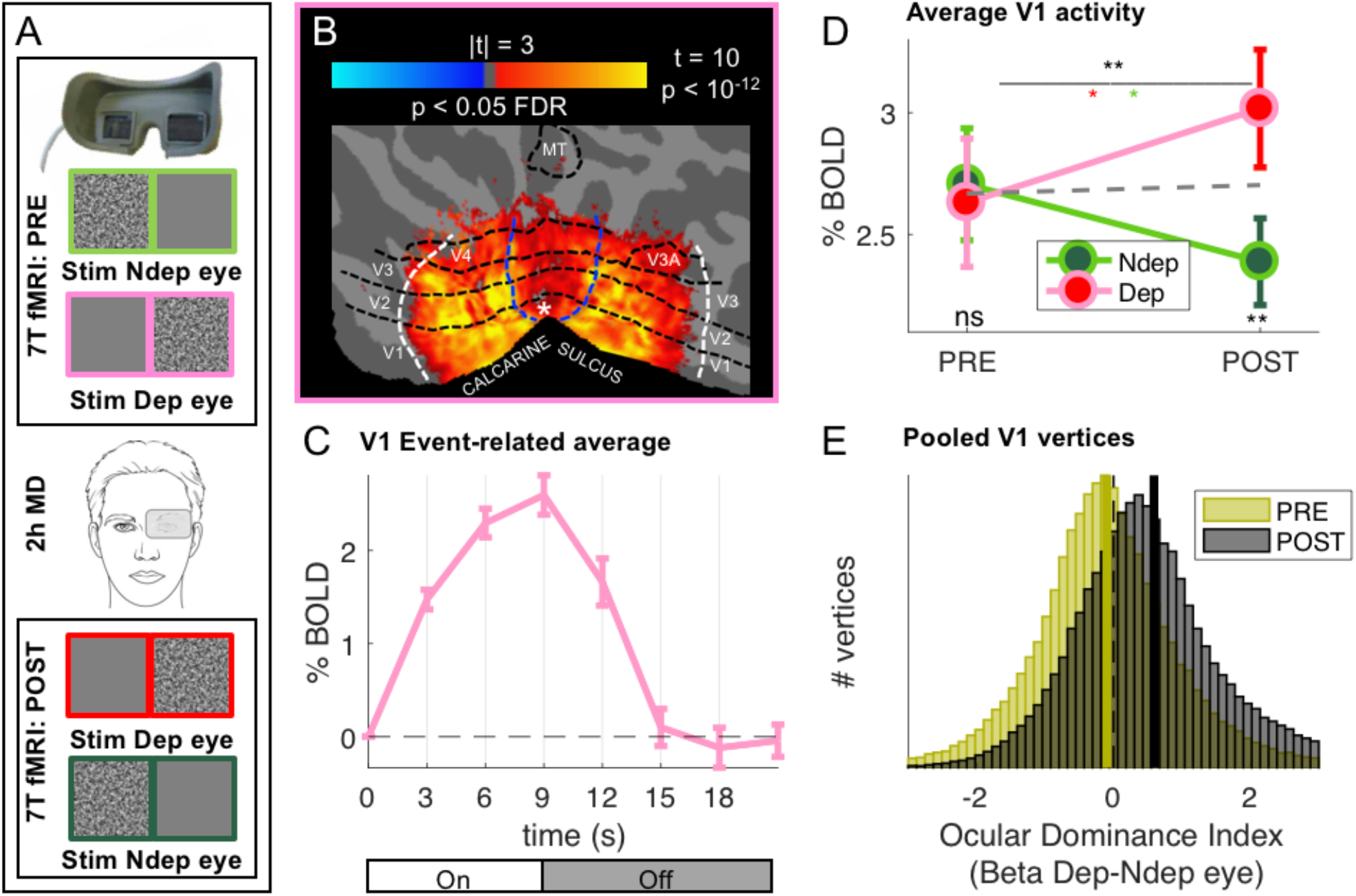
Monocular deprivation modulates 7T BOLD responses in early visual cortex. A: Schematic illustration of the methods. The icons show a band-pass noise stimulus shown to either eye through the MR compatible goggles. Before and after the Pre- and Post-deprivation scans, outside the bore, we also measured binocular rivalry. B: BOLD responses evoked by our band-pass noise stimulus with peak frequency 2.7 cycles per degree (cpd), presented in the deprived eye PRE-deprivation, mapped on the flattened cortical surface, cut at the calcarine sulcus. T-values are obtained by aligning GLM betas for each subject and hemisphere to a left/right symmetric template hemisphere, excluding vertices for which preferred eccentricity was not adequately estimated or smaller than 1 (the same criterion used for al analyses), then evaluating the distribution of betas in each vertex against 0 (one-sample t-test) and FDR correcting across the entire cortical surface. Black dashed lines show the approximate average location of the regions of interested V1 through MT, which were mapped on the individual subject spaces (see methods); white and blue lines represent the outer limits of the representation of our screen space (24 × 32deg) and the foveal representation (< = 1deg, where eccentricity could not be mapped accurately) respectively. C: BOLD modulation during the 3 TRs of stimulus presentation (from 0 to 9s) and the following 4 blank TRs, for the 2.7 cpd noise stimuli delivered to the deprived eye before deprivation. The y-axis show the median percent BOLD signal change in V1 vertices relative to the signal at stimulus onset, averaged across subjects. Error bars give s.e. across participants. Note the small between-subject variability of the response (given that the response of each subject was computed for just two blocks of stimulation-blank). D: Average BOLD response to the band-pass noise stimulus with peak frequency 2.7 cpd, in each of the four conditions, computed by taking the median BOLD response across all V1 vertices then averaging these values across participants (after checking that distributions do not deviate from normality, Jarque-Bera hypothesis test of composite normality, all p > 0.06). The top black star indicates the significance of the ANOVA interaction between factors time (PRE, POST deprivation) and eye (deprived, non-deprived); the other stars report the results of post-hoc t-tests: red and green stars give the significance of the difference POST minus PRE, for the deprived and non-deprived eye respectively; bottom black stars give the significance of the difference deprived minus non-deprived eye before and after deprivation. * p<0.05; ** p < 0.01; *** p < 0.001; ns non-significant. E: Histograms of Ocular Drive Index: the difference between the response (GLM beta) to the deprived and nondeprived eye, computed for each vertex, separately before and after deprivation. Yellow and black lines give the median of the distributions, which are non-normal (logistic) due to excess kurtosis.

The reliability and high signal-to-noise ratio of our system allow us to obtain significant activations of all the main visual areas, with only two blocks of stimulation (Fig. 1B-C show averages across participants, with ROIs limited by stimulus eccentricity, see methods), thereby targeting the first 10 minutes after deprivation, when the perceptual effects are strongest (Lunghi et al., 2011; Lunghi et al., 2013).

We measured the plasticity effect by comparing activity before/after deprivation in response to stimulation in the two eyes with low- and high-spatial frequency bandpass stimuli that differentially stimulate the magno- and parvocellular pathways. Consistent with prior evidence suggesting higher susceptibility to plasticity of the parvocellular pathway (Lunghi, Berchicci, et al., 2015; Lunghi et al., 2011; Lunghi, Emir, et al., 2015; Lunghi & Sale, 2015), we observe a strong effect of Monocular Deprivation on BOLD responses to stimuli of high spatial frequency (peak 2.7 cycles per degree, high-frequency cut-off at half-height 7.5 cpd). Fig. 1D shows that the V1 response to the high spatial frequency stimuli presented in the left and right eye is nearly equal before deprivation (“PRE”). However, after deprivation (“POST”), the response in the two eye changes in opposite directions, with a boost of the BOLD response (measured as GLM Beta values, expressed in units of % signal change) of the deprived eye and a suppression of the non-deprived eye (see also supplementary Fig. S1). This was formally tested with a two-way repeated measure ANOVA, entered with the mean BOLD responses across all vertices in the left and right V1 region, for the four conditions and each participant (Fig. 1D show averages of this values across participants). The result reveals a significant interaction between the factors *time* (PRE, POST deprivation) and *eye* (deprived, non-deprived; interaction term F(1,18) = 13.80703, p = 0.00158; the result survives a split-half reliability test: see supplementary Fig. S2).

Fig. 1E confirms these findings with an analysis of the aggregate subject data, obtained by pooling all V1 vertices across all subjects. For each vertex, we defined an index of Ocular Dominance computed as the difference of BOLD response to the deprived and non-deprived eye (not to be confused with the anatomical arrangement of vertices with different eye preference that define the ocular dominance columns (Cheng, Waggoner, & Tanaka, 2001; Yacoub, Shmuel, Logothetis, & Ugurbil, 2007)). Before deprivation, the Ocular Dominance index is symmetrically distributed around zero, indicating a balanced representation of the two eyes before deprivation (yellow distribution in Fig.1E). After deprivation (black distribution in Fig.1E), the Ocular Dominance distribution shifts to the right of 0, indicating a preference for the deprived eye (non-parametric Wilcoxon sign-rank test comparing the PRE and POST Ocular Dominance medians, z = 115.39, p < 0.001).

In principle, the boost of responses to the deprived eye seen in Fig. 1D could be produced by enhancing the response of vertices that originally preferred the deprived eye (without shifting ocular dominance) or by changing Ocular Dominance of vertices that originally preferred the non-deprived eye, driving them to prefer the deprived eye. The shift of the Ocular Dominance histogram in Fig. 1E is more compatible with the latter case, implying a recruitment of cortical resources for the representation of the deprived eye. To investigate this further, we monitored the final POST-deprivation Ocular Dominance of individual vertices that, PRE-deprivation, preferred the deprived eye (yellow half distribution in Fig 2A). The majority of vertices continue to prefer the same eye before and after deprivation. The median Ocular Dominance is significantly larger than 0 both PRE and POST (Wilcoxon sign-rank test, z > 101.54, p < 0.0001 in both cases) and the correlation between Ocular Dominance indices before and after deprivation is strong and positive (Pearson’s R(32236) = 0.22 [0.21-0.23], p < 0.0001). Note that a completely random reassignment of Ocular Dominance after deprivation would have produced a histogram centered at 0 and no correlation between Ocular Dominance indices PRE- and POST deprivation. This is not consistent with the results of Fig. 2B, which thereby provide evidence that our estimates of Ocular Dominance before and after deprivation are congruent, even though they were collected in different fMRI sessions separated by 2h. In addition, the distribution of Ocular Dominance after deprivation is well predicted by adding only a small amount of noise to the original half distribution (Gaussian noise with 0.12 standard deviation, black line), suggesting that these vertices were largely unaffected by monocular deprivation. This is also supported by the repeated measure ANOVA of individual subject data (Fig. 2A), revealing a strong main effect of eye (F(1,18) = 48.28901, p < 10^−5^): the response to the deprived eye is stronger than the non-deprived eye, both before deprivation (due the selection, t(18) = −8.616, p < 10^−5^), and after deprivation (t(18) = −4.281, p < 10^−5^), with no effect of time and no *time × eye* interaction (all F(1,18) = 0.20429, p > 0.5).

**Figure 2:**
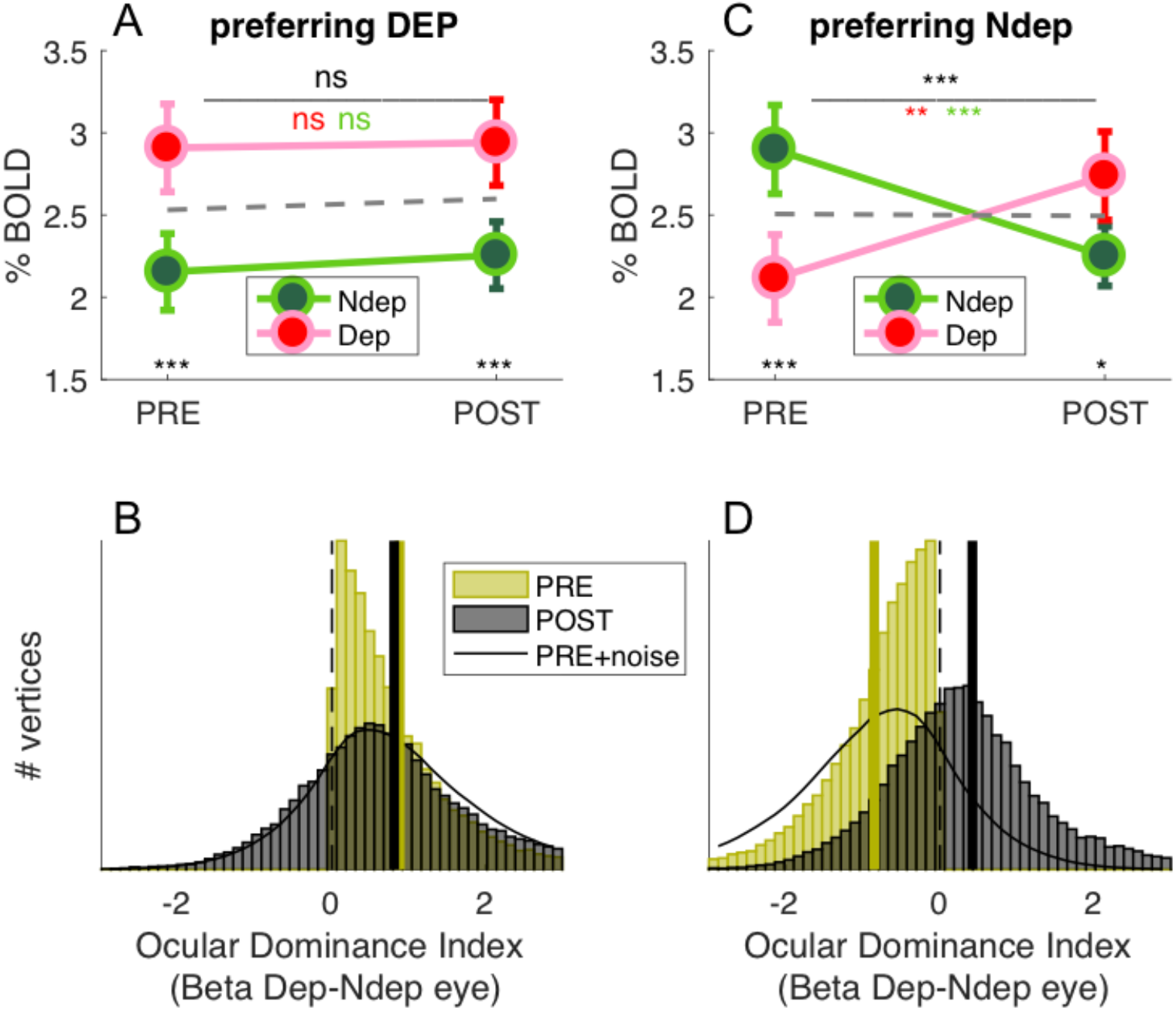
Monocular deprivation shifts 7T BOLD Ocular Dominance in V1. A & C: Average BOLD responses with the same conventions as in Fig. 1D but analysing data from two subregions of V1. A: only vertices that, before deprivation, respond preferentially to the deprived eye. C: only vertices that, before deprivation, respond preferentially to the non-deprived eye. B & D: Histograms of Ocular Dominance Index (as for Fig. 1E), in the two sub-regions of V1, computed before and after deprivation. The black curve simulates the result of adding random noise to the distribution obtained before deprivation; only in B does this approximate the distribution observed after deprivation.

A completely different pattern is observed for the vertices originally preferring the non-deprived (yellow half-distribution in Fig. 2D). Here the distribution of Ocular Dominance clearly shifts after deprivation; the median moves from significantly negative before deprivation (Wilcoxon sign-rank test, z = −175.97, p < 0.0001) to significantly positive after deprivation (Wilcoxon sign-rank test, z = 64.46, p < 0.0001), implying a shift of dominance in favor of the deprived eye. Again, this is not consistent with a random reassignment of Ocular Dominance after deprivation, which predicts a distribution centered at 0. Contrary to Fig. 2B, the POST-Ocular Dominance distribution cannot be predicted by injecting Gaussian noise to the PRE-Ocular Dominance distribution (black line, 0.12 standard deviation like for Fig. 2B): for these vertices, there is a shift of Ocular Dominance with short term monocular deprivation. This is confirmed with the repeated measure ANOVA (Fig. 2C), where the *time × eye* interaction is significant (F(1,18) = 44.82812, p < 10^−5^), implying a different modulation PRE and POST deprivation. In addition and crucially, POST-deprivation BOLD responses to the deprived eye are significantly larger than POST-deprivation responses to the non deprived eye (t(18) = −2.775 p = 0.012; whereas, by selection, the opposite is true before deprivation: t(18) = 12.034, p < 10^−5^).

In summary, Ocular Dominance before deprivation defines two similarly sized sub-regions of V1 vertices (44.58 ± 5.38% and 55.42 ± 5.38% of analyzed V1 vertices; 44.84 ± 5.12% and 55.16 ± 5.12% of all V1 vertices) with radically different behaviors that are not consistent with an artifact induced by vertex selection. The sub-region that originally represents the deprived eye does not change with deprivation; the sub-region that originally represents the other non-deprived is rearranged with deprivation, as a large portion of vertices turn to prefer the deprived eye.

If plasticity were not eye-specific and/or we failed to match our V1 vertices before/after deprivation, we would expect that splitting the distribution of V1 ocular dominance generates opposite effects in the two subpopulations: vertices preferring the deprived eye before deprivation should swap to prefer the other eye, mirroring the effect seen in the vertices preferring non-deprived eye. This is not seen, implying that we did successfully match vertices across the 2h of deprivation and that the selective Ocular Dominance shift, observed for about half of our vertices, is not an artifact.

We also measured the perceptual effects of short-term monocular deprivation affects using Binocular Rivalry, just before the PRE- and POST-deprivation fMRI sessions. In line with previous studies (Binda & Lunghi, 2017; Lunghi, Berchicci, et al., 2015; Lunghi et al., 2011; Lunghi, Emir, et al., 2015; Lunghi & Sale, 2015), short-term monocular contrast deprivation induced a 30% increase of phase duration for the deprived eye (POST to PRE-deprivation ratio: 1.31 ± 0.30) and a 15% decrease of phase duration for the non-deprived eye (ratio: 0.86 ± 0.30), producing a significant *time × eye* interaction (Fig. 3A, repeated measure ANOVA on the mean phase durations for each participant, interaction: F(1,18) = 23.56957, p = 0.00013). We defined a psychophysical index of the deprivation effect (DI_psycho_) by using Eq. 6 in methods section, where the POST to PRE-deprivation ratio of phase durations for the deprived eye, is divided by the same ratio for the non-deprived eye. Values larger than 1 imply a relative increase of the deprived eye phase duration, i.e. the expected effect; a value less than 1 indicates the opposite effect and a value of 1 indicates no change of mean phase duration across eyes. All but two subjects have values larger than 1, indicating a strong effect of deprivation. However, the scatter is large with values ranging from 0.7 to 3, suggesting that susceptibility to visual plasticity varies largely in our pool of participants.

**Figure 3:**
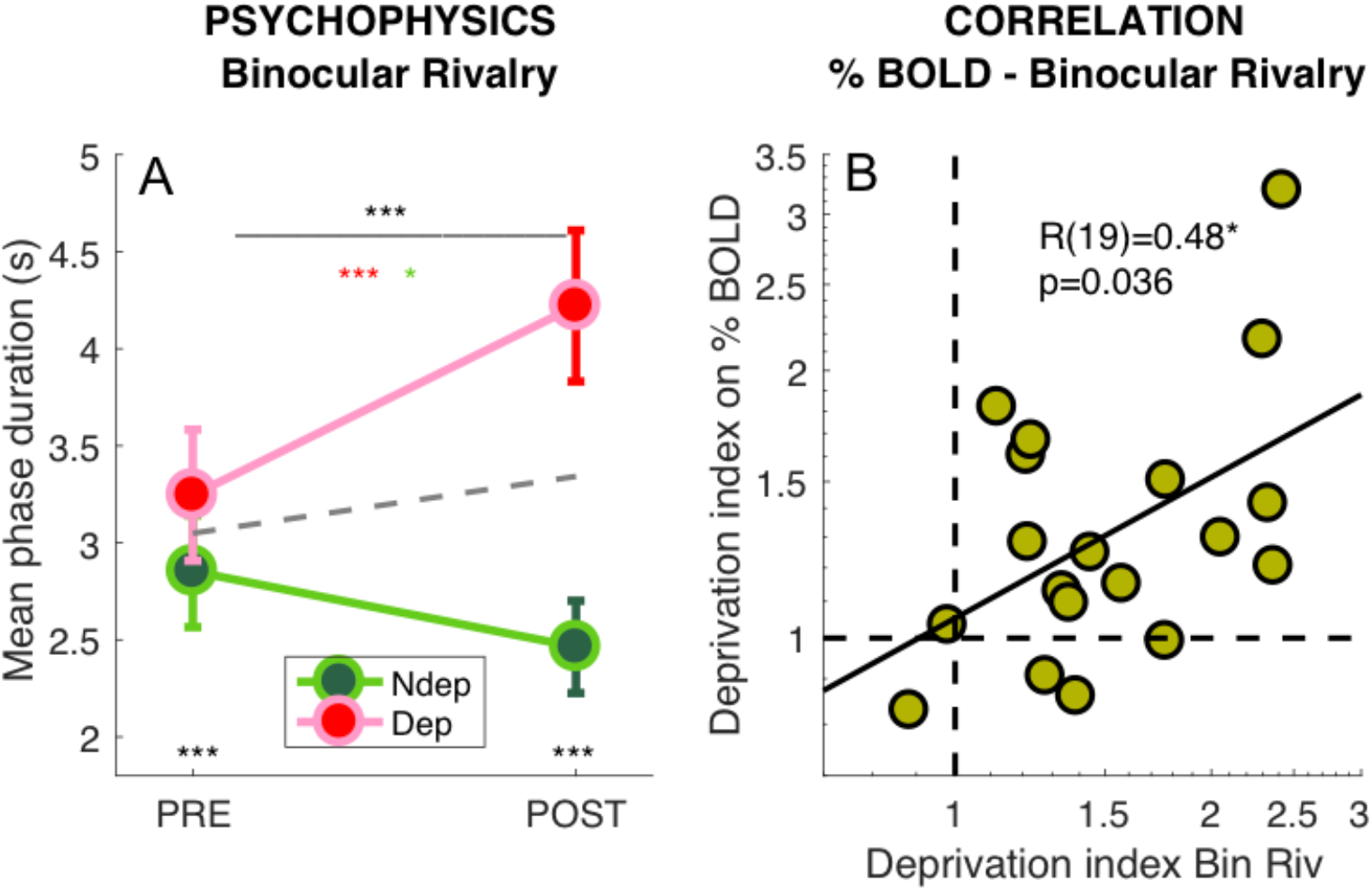
Deprivation effects on BOLD and on psychophysics are correlated. A: Effect of deprivation on Binocular Rivalry dynamics. Average phase duration for the deprived and nondeprived eye, before and after deprivation, same conventions as in Fig. 1D. Mean phase duration distributions do not deviate from normality (Jarque-Bera hypothesis test of composite normality, all p > 0.171) B: Correlation between the deprivation index (the POST to PRE-ratio for the deprived eye divided by the same ratio for the non-deprived eye, Eq. 6 in Methods) computed for the binocular rivalry mean phase duration and for the BOLD response to our band-pass noise stimulus with peak frequency 2.7 cpd. Text insets show the Pearson’s correlation coefficient and associated p-value.

Capitalizing on this variability, we tested whether the size of the psychophysical effect correlates with the BOLD effect across participants. Using the same Eq. 6 to compute the deprivation effect on BOLD responses (DI_BOLD_), we observed a strong correlation between the effect of monocular deprivation on psychophysics and BOLD (shown in Fig. 3B). Subjects who showed a strong deprivation effect at psychophysics (DI_psycho_ > 2) also showed a strong deprivation effect in BOLD responses (DI_BOLD_ = 1.85 ± 0.42). Given that the psychophysics was measured only for central vision and at 2 cpd stationary grating, whereas BOLD responses were pooled across a large portion on V1 and were elicited using broadband dynamic stimuli, the correlation suggests that the psychophysical effect may be used as a reliable proxy of a general change of cortical excitability, which can be measured by fMRI.

### Monocular deprivation shifts BOLD Spatial Frequency Tuning for the deprived eye

The BOLD measure we use here gives us the chance to measure the effect of Monocular Deprivation across spatial frequencies and as function of eccentricity. We used 5 band-pass noise (1.25 octaves half-width at half maximum) stimuli with peak spatial frequency selected to have a complete coverage of spatial frequencies from 0.03 to 12.5 cpd (see Supplementary Fig. S3). In contrast with the strong and reliable plasticity of responses to the high spatial frequency stimulus (peaking at 2.7 cpd, Figs. 1–3), we find that the plasticity effect is absent at low spatial frequencies (interaction index for the highest spatial frequency stimulus: 0.70±0.19; for the lowest spatial frequency stimulus: 0.16±0.15; paired t-test t(18) = −3.441, p = 0.003).

Thus, monocular deprivation produces a change of the spatial frequency selectivity of the V1 BOLD response. Before deprivation, the BOLD response shows a broad band-pass selectivity for our stimuli, with a preference for the stimulus peaking around 1 cpd, and a slight attenuation at higher spatial frequencies, similar for the two eyes (Fig. 4A). After deprivation (Fig. 4B), the non-deprived eye shows similar selectivity and an overall decrease of responses. For the deprived eye, the shape of the curve changes: from band-pass to high-pass, implying that the enhancement affects primarily the higher spatial frequencies.

**Figure 4:**
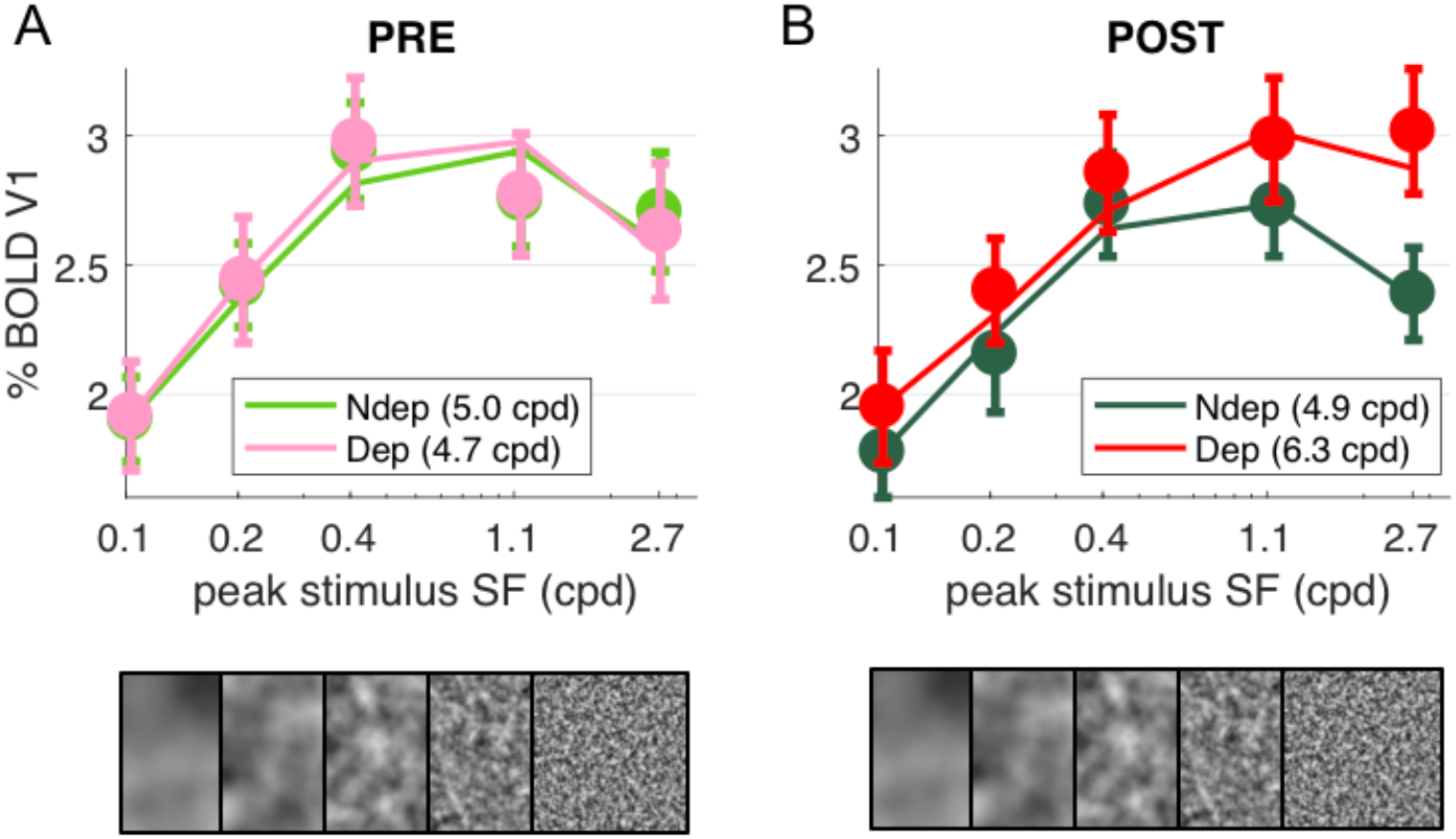
Deprivation affects spatial frequency selectivity in V1. V1 BOLD responses to all five of our band-pass noise stimuli (with peaks at 0.1, 0.2, 0.4, 1.1 and 2.7 cpd, see spectra in supplementary Fig. S2); A: response to stimuli in either eye, before deprivation; B: response to stimuli in either eye, after deprivation. Responses are computed as medians across all V1 vertices (like in Fig. 1D), averaged across subjects (error bars report s.e.m.). Continuous lines show the response of the best-fit population Spatial Frequency tuning (with the one parameter, the high spatial frequency cut-off, indicated in the legend), estimated by applying to the average V1 BOLD response the same model used to predict individual vertex responses fitting procedure illustrated in supplementary Fig. S3).

To model this effect, we assume that each vertex on the cortical surface subtends a multitude of neuronal channels, each with narrow tuning for spatial frequency and collectively spanning a large range of spatial frequencies – an approach conceptually similar to the population Receptive Field model for retinotopic mapping (Dumoulin & Wandell, 2008). Independently of the exact bandwidth and peak preference of the neuronal population contributing to the final BOLD selectivity, we find that the shape of all these curves is captured with a simple one-parameter model: what we term the population tuning for Spatial Frequency. This is given by a Difference-of-Gaussians (DoG) function with one free parameter, the spatial constant (while the excitation/inhibition spatial constant ratio is fixed; see eq. 4 in the Methods and curves in Supplementary Fig. S4). The free parameter sets the high spatial frequency cut-off at half-height of the filter. The continuous lines in Fig. 4 show how the model fits the grand-average of V1 responses, with best fit cut-off around 5 cpd similar for all conditions except for the POST-deprivation deprived eye, where the cut-off is 6.2 cpd (single vertex examples are given in supplementary Fig. S4C-I). The DoG equation has been successfully used in previous studies to model CSF and neural responses at variable stimulus parameters e.g. illumination levels (Enroth-Cugell & Robson, 1966; Hawken, Parker, & Lund, 1988), validating this equation for modeling the overall selectivity of large neuronal ensembles.

Using this model to analyze single vertex responses, we evaluated the best-fit spatial frequency cut-off of the neural population contributing to the vertex BOLD response (see details in the methods and Supplementary Fig. S4A-C; briefly, we used the DoG model to predict the response elicited by our five band-pass noise stimuli in populations with different spatial frequency selectivity, i.e. filters with different cut-off; we then found the cut-off value that maximizes the correlation between the predicted responses and the observed BOLD responses). We used this procedure to fit BOLD responses in each of our four conditions, estimating spatial frequency selectivity in individual vertices in each condition: separately for the two eyes, PRE/POST deprivation. Before deprivation, the spatial frequency cut-off decays with eccentricity as expected. Fig 5A maps both eccentricity (pRF eccentricity estimates from a separate retinotopic mapping scan) and spatial frequency cut-off values, obtained by fitting responses to the deprived eye, before deprivation (averaged across hemispheres and subjects). The cut-off is around 16 in the para-fovea (eccentricity around 1.5 deg) and down to 4 in the periphery (eccentricity around 8 deg). This relationship between eccentricity and spatial frequency preference is consistent with previous fMRI results (D’Souza, Auer, Frahm, Strasburger, & Lee, 2016; Henriksson, Nurminen, Hyvarinen, & Vanni, 2008) and with psychophysics (Rovamo, Virsu, & Nasanen, 1978). The model captures well the selectivity of an example V1 vertex (Fig. 5B, goodness of fit better than 0.9), sampled from the mid-periphery (3.4 deg) for the deprived eye, both before and after deprivation. The spatial frequency cut-off after deprivation shifts to higher values, increasing (in this example) by about a factor of three. Fig. 5C-D shows that this behavior is systematically observed across V1 vertices, but only for the deprived eye. Here the average cut-off is plotted as function of eccentricity, and the roll-off is consistent with the map in Fig. 5A. For the non-deprived eye, there is no effect of deprivation on spatial frequency selectivity (Fig. 5C). In contrast, for the deprived eye (Fig. 5D), there is a shift towards preferring higher spatial frequencies, at all eccentricities, which is captured by an increased value of the cut-off frequency parameter leading to an increased acuity of the BOLD response to the deprived eye.

**Figure 5:**
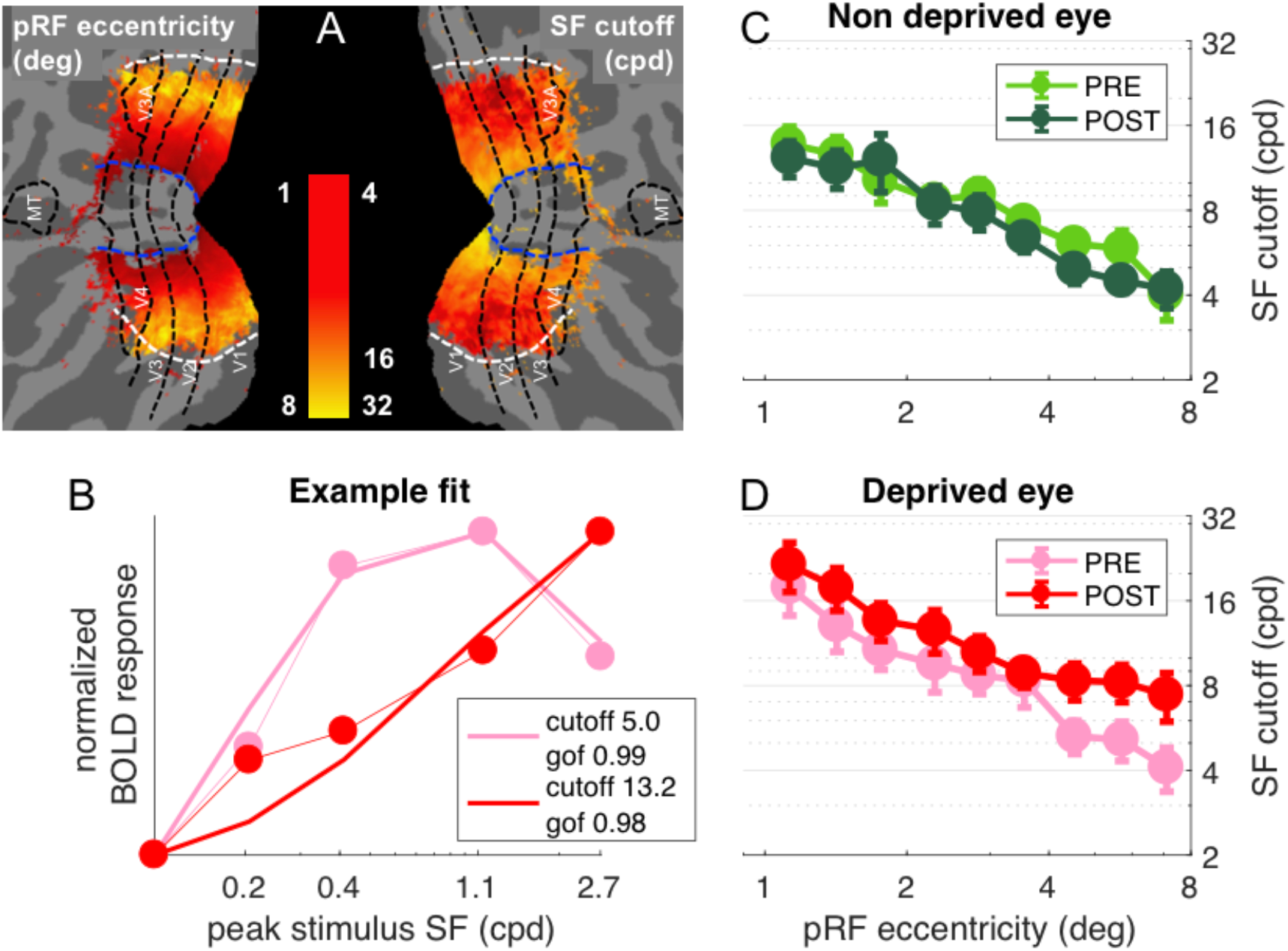
population Spatial Frequency Tuning in V1. A: Maps of pRF eccentricity and best fit spatial frequency cut off (for the deprived eye before deprivation) after aligning the parameter estimates for all hemispheres to a common template and averaging them across subjects and hemispheres, after excluding vertices for which the average preferred eccentricity was not adequately estimated or smaller than 1 (the same exclusion criteria used for analyses). B: Predicted and observed BOLD activity in one example vertex, elicited in response to our bandpass noise stimuli in the deprived eye PRE (pink) and POST deprivation (red), with best fit spatial frequency cut off (reported in the legend). C-D: Best fit spatial frequency cut-off, averaged in sub-regions of V1 defined by pRF eccentricity bands, and estimated separately for the two eyes and PRE/POST deprivation.

Note that the change of spatial frequency selectivity for the deprived eye is most evident at eccentricities of 4 deg and higher (see Fig. 5D), where vertices have peak sensitivity at mid-to-low spatial frequencies before deprivation. In the fovea, where many vertices already prefer the highest spatial frequency stimulus before deprivation, our fitting procedure is likely to underestimate the change of spatial frequency selectivity. Importantly, the spatial frequency selectivity for the non-deprived eye is unchanged at all eccentricities, corroborating the eye and stimulus-specificity of the short-term monocular deprivation effect.

To test the significance of these effects, we pooled the best fit cut-off values from all selected V1 vertices across eccentricities and averaged them across participants (Fig. 6A). The repeated measure ANOVA (performed on the log-transformed values, which are distributed normally as assessed by the Jarque-Bera test) shows no significant *time × eye* interaction (F(1,18) = 3.67607, p = 0.07121) and non significant main effect of time (F(1,18) = 2.62546, p = 0.12255) but a significant main effect of eye (F(1,18) = 13.58079, p = 0.00169). This is clarified by post-hoc t-tests revealing that the increase of spatial frequency cut-off for the deprived eye is significant (t(18) = −2.263, p = 0.036) whereas there is no significant change for the non-deprived eye (t(18) = 0.440, p = 0.665). Given that the *time × eye* interaction in the full V1 region is not significant, and to minimize noise contamination, we evaluated the effect of deprivation on spatial frequency cut-off at the individual level by a “Deprived Eye Change (DepC_cutoff_)” index (Eq.7 in the methods), i.e. taking the POST vs. PRE-deprivation ratio of the spatial frequency cut-off for the deprived eye alone. As this ratio varies widely across participants, over more than 3 octaves, we asked whether this variability correlates with our psychophysical probe of plasticity: binocular rivalry. We used the same Eq. 7 to index the psychophysical change of the deprived eye (DepC_psycho_), the POST to PRE-ratio of mean phase duration for the deprived eye, and found a strong positive correlation (Fig. 6B). POST-deprivation, the deprived eye shows an increase of mean phase duration (in binocular rivalry) and an increase of the spatial frequency cut-off (best fit of the BOLD responses): participants showing a stronger increase of phase duration, also showed a larger shift of selectivity towards higher spatial frequency. The correlation is consistent with the result of Fig. 3 showing that the enhancement of BOLD responses is correlated with the change of binocular rivalry and selective for the highest spatial frequency stimulus.

**Figure 6:**
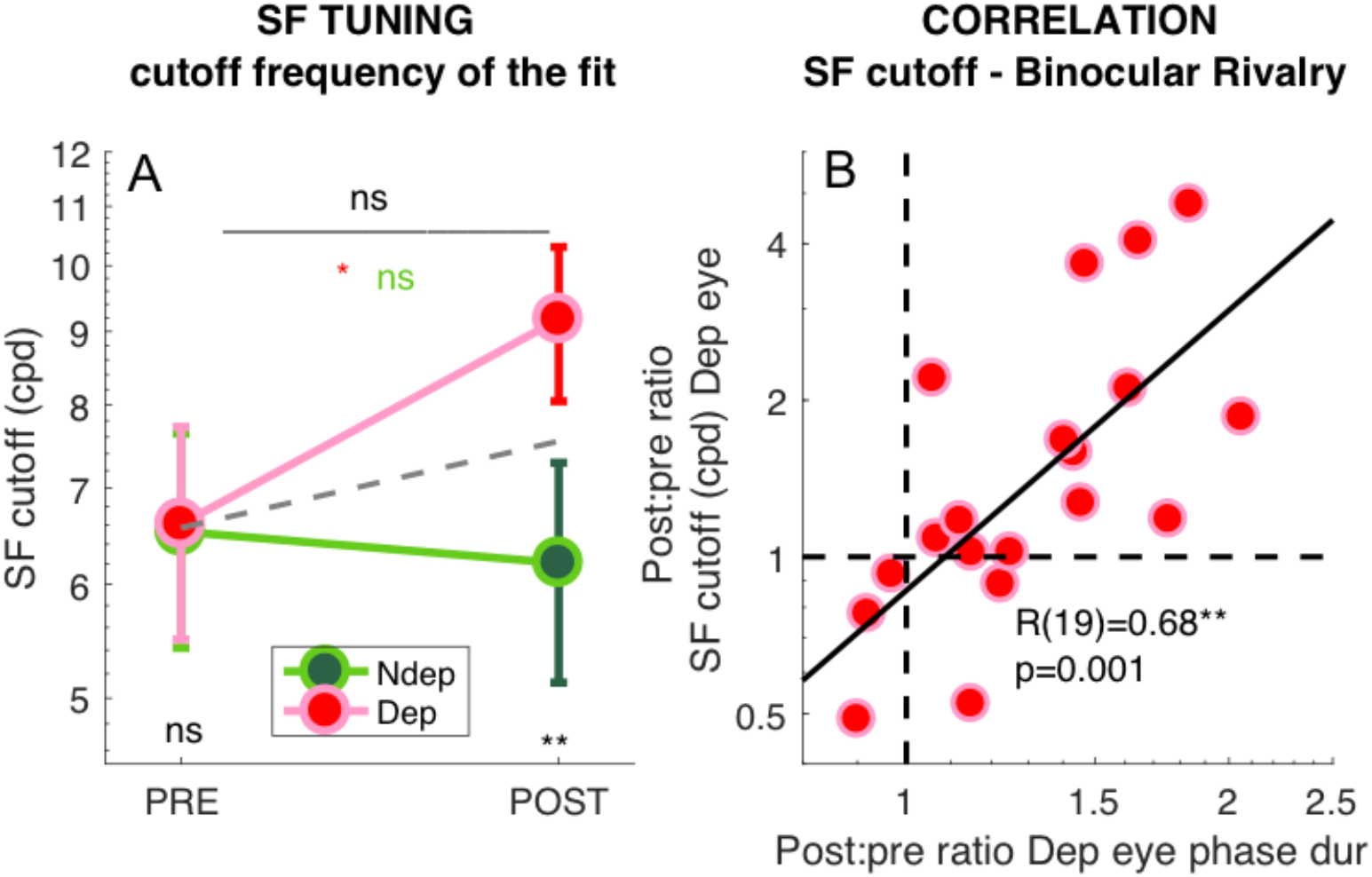
Deprivation effects on the deprived eye population Spatial Frequency Tuning and binocular rivalry phase duration are correlated. A: Effect of deprivation on spatial frequency cut off values. Average cut-off across all V1 vertices (pooled across eccentricities) for the deprived and non-deprived eye, before and after deprivation, same conventions as in Fig. 1D. Distributions of the log-values do not deviate from normality (Jarque-Bera hypothesis test of composite normality, all p > 0.285). B: Correlation between the POST/PRE ratio (Eq. 7 in the Methods) computed for the binocular rivalry mean phase duration and for the spatial frequency cut off for the deprived eye.

### Monocular deprivation affects BOLD responses in the ventral stream areas beyond V1

We measured the effect over the main extra-striate visual cortical areas. The selective boost of the deprived eye response to the high spatial frequency is as strong in V2 as in V1 (Fig. S1 and Fig. 7E). The boost is present also in V3 and V4. In V4 the boost appears to be present also for lower spatial frequencies, but again only for the deprived eye (Fig. 7A-B), possibly reflecting the larger spatial frequency bandwidth of V4 neurons compared to V1.

**Figure 7:**
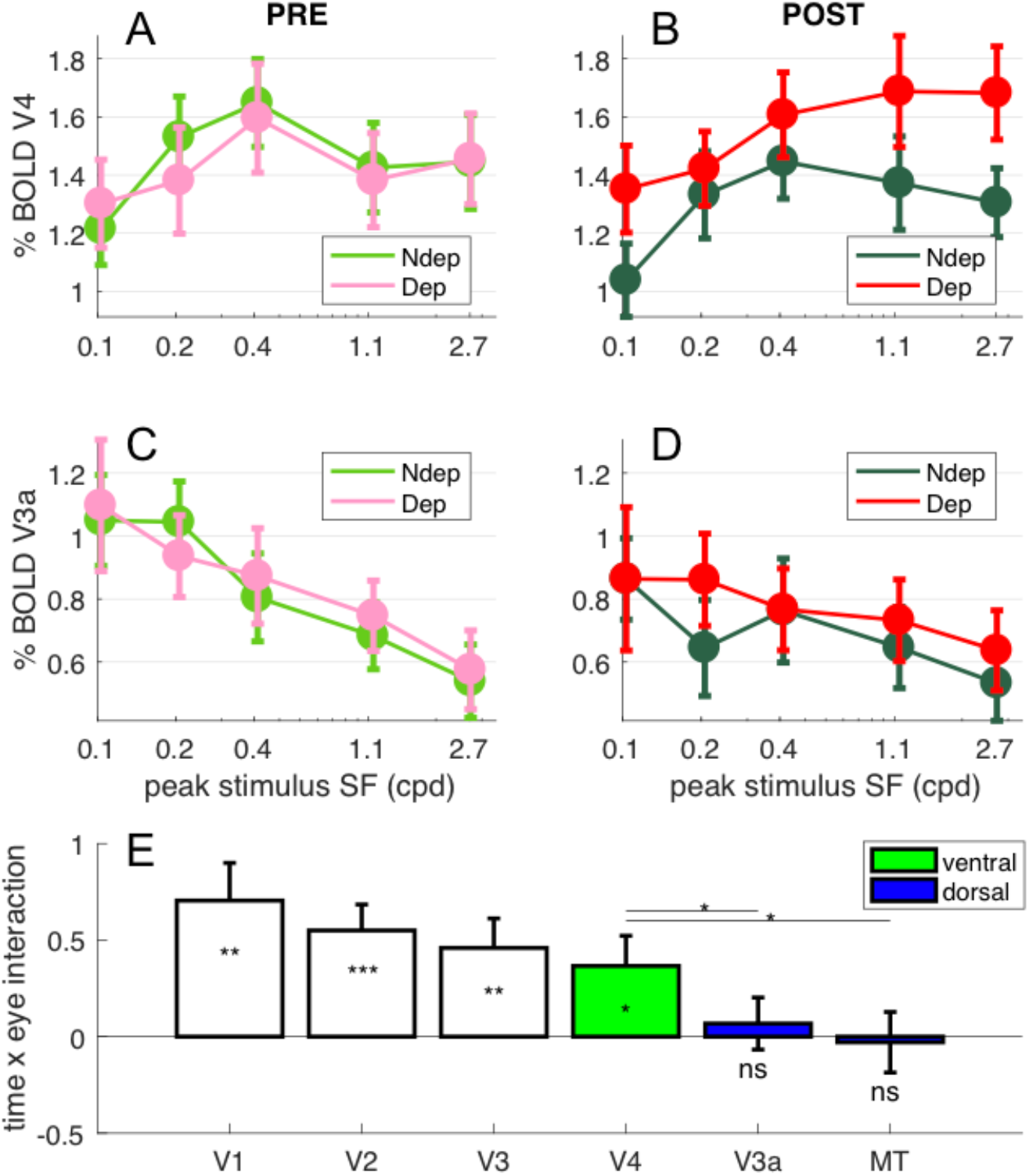
Deprivation effects are stronger in ventral than in dorsal stream areas. Panels A-B show V4 responses across spatial frequency stimuli presented to each eye (colored lines) before (A) and after deprivation and panels C-D show V3a responses. Each data point is computed by taking the median BOLD response across vertices in the region of interest for each stimulus and subject, then averaging across subjects (errorbar report s.e.m.). Panel E summarizes the effect of deprivation measured for the highest spatial frequency stimulus in the V1, V2, V3/VP, V4, V3a and MT region of interest, computing the interaction term (POST-PRE difference of BOLD response for the deprived eye, minus the same value for the non-deprived eye) for individual participants and the 2.7 cpd stimulus. Values around 0 indicate no effect of deprivation and values larger than 0 indicate a boost of the deprived eye after deprivation. One-sample t-tests comparing this value against 0 give a p-value equivalent to that associated with the interaction term of the ANOVA (Fig. 1D); the significance of the resulting t-value is given by the stars plotted below each errorbar. Stars plotted above the lines show the results of paired t-tests comparing interaction terms in V4 and V3a/MT. *** = p<0.001; ** = p<0.01; * = p<0.05; ns = p>=0.05. Green and Blue highlight the assignment of the higher tier areas to the ventral and dorsal stream respectively.

The results are very different for dorsal area V3a (Fig. 7C-D), where there is no significant change of responses in either eye. Moreover, the V3a preference moves to lower spatial frequencies, consistent with a stronger input of the magnocellular pathway to the dorsal visual stream (Henriksson et al., 2008; Singh, Smith, & Greenlee, 2000).

Fig. 7E quantifies the effect of short-term monocular deprivation (using the ANOVA time x eye interaction term, which measures the eye-selective modulation of BOLD response after deprivation for the highest spatial frequency) across the main visual areas. The plasticity effect is strongest in V1, V2 and V3; it is still strong and significant in ventral area V4 (t(18) = 2.41 p = 0.0270), but it is absent in V3a and MT, where the time x eye interaction is not significantly different from 0 (t(18) = 0.52 p = 0.6115 and t(18) = −0.19 p = 0.8513 respectively). The plasticity effect in ventral area V4 is significantly stronger than in dorsal areas V3a and MT (t(18) = 2.39, p = 0.0278 and t(18) = 2.36, p = 0.0299 for V4-V3a and V4-MT respectively).

This result suggests a preferential involvement of the parvocellular vs. magnocellular pathway, leading to the differential plasticity effect in extra-striate visual areas of the ventral and dorsal pathway. Interestingly, the plasticity effect is robust in areas where the majority of cells are binocular (like V3 and V4), indicating that the effect does not require segregated representations of the two eyes (e.g. ocular dominance columns).

## Discussion

We demonstrate that two hours of abnormal visual experience has a profound impact on the neural sensitivity and selectivity of V1. BOLD activity across the V1 cortical region paradoxically increases for the eye that was deprived of contrast vision, with the strongest boost for the higher spatial frequency stimuli, and decreases for the eye exposed to normal visual experience.

The enhanced response to the deprived eye contrasts with the established weakening of the deprived eye response after a brief deprivation during the critical period (Gordon & Stryker, 1996; Kiorpes et al., 1998; Levi & Carkeet, 1993; Wiesel & Hubel, 1963) and a long-term deprivation in adulthood (Sato & Stryker, 2008). However, this fits well with the concept of homeostatic plasticity, the tendency of neural circuits to keep the average firing rates constant in spite of anomalous stimulation. First observed in animal models deprived for few days during the critical period, homeostatic plasticity induces a transient boost of the deprived eye, mediated by a modulation of the response gain (Mrsic-Flogel et al., 2007; G. G. Turrigiano & Nelson, 2004). More recently, similar observations have been made in the adult macaque V1 after two hours of monocular deprivation during anesthesia (Fregnac et al., 1988). The post-deprivation gain boost observed in the monkey is consistent with our observations of an increased BOLD response to the deprived eye. We also observe an antagonistic suppression of the non-deprived eye BOLD response; together, the two effects lead to a shift of ocular preference of individual vertices in favor of the deprived eye. However, this effect is only observed in those V1 vertices that, before deprivation, responded preferentially to the non-deprived eye. No such change of ocular preference is seen in vertices that already prefer the deprived eye before deprivation, which maintain their eye-preference after deprivation. This pattern of results cannot be explained by any overall gain increase; rather, it is consistent with the idea that the representation of the deprived eye recruits cortical resources (which may or may not correspond to cortical territory) normally dedicated to the other eye.

A similar antagonist effect on the two eyes (boosting the deprived eye and suppressing the non-deprived eye) was also observed in the VEP responses after short-term monocular deprivation (Lunghi, Berchicci, et al., 2015) and could be implemented through a modulation of the excitatory/inhibitory circuitry. Regulation of the excitation/inhibition balance through GABAergic signaling is considered to be a key factor for cortical plasticity, including homeostatic plasticity (Maffei & Turrigiano, 2008). Interestingly, the involvement of GABAergic signaling in the effect of short-term monocular deprivation is directly supported by MR Spectroscopy data in adult humans, showing that resting GABA in a large region of the occipital cortex is specifically reduced after short-term monocular deprivation (Lunghi, Emir, et al., 2015).

The functional relevance of the BOLD changes we observe is demonstrated by their correlation with our behavioral assay of plasticity, obtained through binocular rivalry. This correlates both with the BOLD ocular dominance change (relative boost/suppression of the deprived/non-deprived eye), and with the BOLD acuity change for the deprived eye (change of spatial frequency tuning, assessed with our pRF-like modeling approach). The correlation holds despite binocular rivalry being restricted to foveal vision, whereas the assessment of BOLD plasticity is pooled across V1 (including the mid-periphery). This implies that the change of binocular rivalry dynamics is a proxy for the more general plasticity effects that involves the whole primary visual cortex. This finding has long reaching implications, as it could validate the use of binocular rivalry as a biomarker of adult cortical plasticity, based on the neural mechanisms revealed by the present 7T fMRI results. Interestingly, the binocular rivalry phenomenon originates in the primary visual cortex – probably at the earliest stages – and is an expression of the dynamics of excitatory transmission and inhibitory feedback (Tong, Meng, & Blake, 2006); as such it is a measure that could reflect the overall excitation-inhibition ratio (van Loon et al., 2013), and its modulation in plasticity (Lunghi, Emir, et al., 2015; Maffei & Turrigiano, 2008).

Other evidence show that selective deprivation of orientation (Zhang et al., 2009) or spatial frequency (Zhou et al., 2014) or color (Zhou, Reynaud, Kim, Mullen, & Hess, 2017) may lead to a boost of the deprived signal. These effects have been interpreted as a form of release of inhibition from the adapted signal (Zhang et al., 2009) – a concept that is not distant from homeostatic plasticity, where the network aims to keep overall activity constant. The conceptual border between adaptation and plasticity is fussy, given that some mechanisms are shared and both effects have the same outcomes. Be it adaptation or plasticity, the monocular deprivation mechanisms are probably cortical and affect mainly the deprived eye. There is evidence that the boost of the deprived eye is also observed when the two eyes receive equally strong stimulation, but perception of one eye stimulus is suppressed experimentally (by the continuous flash suppression technique Kim, Kim, & Blake, 2017); this result dismiss the retinal or thalamic contribution to the deprivation effect. Only in rare occasion adaptation induce effect that last over days, yet our recent work shows that deprivation effects of short-term monocular deprivation is retained across 6h sleep (Menicucci et al., 2018), consistent with plasticity reinforcement during sleep (Raven, Van der Zee, Meerlo, & Havekes, 2018; Timofeev & Chauvette, 2017). Most importantly, in adult amblyopic patients, short-term monocular deprivation is able to induce improvement of visual acuity and stereovision (Lunghi et al., 2016) for more than one year. All this evidence supports the concept that short-term plasticity is possible in the human adult cortex and corroborate more direct evidence of functional plastic changes provided by short-term paired TMS studies (Chao et al., 2015) – which, interestingly, produce transient changes that decay with a similar time-course as the monocular deprivation effect, within about our hour. Also, Hebbian changes at the single cell level can be observed in V1 of adult anaesthetized cat, following activity pairing over a similar time-scale (from minutes to a few hours) (Fregnac et al., 1988), clearly demonstrating the plasticity potential of the adult V1.

Given these strong indications that the effect of monocular deprivation is a form of plasticity, our observation that the effect is present in V1 calls for a reconsideration of the established idea that human adult primary sensory cortex is resilient to plastic change. The perceptual learning literature suggest that associative cortical areas, but not V1, retain a high degree of flexibility (B. Dosher & Lu, 2017; B. A. Dosher & Lu, 1999; Fuchs & Flugge, 2014; Harris, Gliksberg, & Sagi, 2012; Kahnt, Grueschow, Speck, & Haynes, 2011; A. Karni et al., 1995; Lewis, Baldassarre, Committeri, Romani, & Corbetta, 2009; Shibata et al., 2012; Watanabe & Sasaki, 2015); our data support the original notion that V1 circuitry may be optimized by perceptual experience (Fiorentini & Berardi, 1980). However, we do observe a plasticity effect beyond V1. This is strong in V2 and V3; after V3, a clear difference emerges between extra-striate visual areas in the ventral and dorsal stream. While ventral area V4 still shows a strong plasticity effect, area V3a, located at a similar tier in the dorsal stream, shows no modulation with short-term monocular deprivation. V4 is a primary target of the parvocellular system, which is best stimulated by our highest spatial frequency stimulus; V3a and MT are preferential targets of the magnocellular system, which respond more strongly to our lower spatial frequency stimuli (see Fig. 7). The different plasticity response of the ventral and dorsal stream, together with the selectivity for the high spatial frequencies of the V1 plasticity, suggests that the parvocellular pathway is most strongly affected by short-term plasticity. This fits well with several sources indicating that selective deprivation of the stimuli that optimally drive the parvocellular system is sufficient to produce a reliable plasticity effect (Begum & Tso, 2016). It is consistent with the finding that the effect of short-term monocular deprivation is strongest and most durable when tested with chromatic equiluminant stimuli (Lunghi et al., 2013).

Understanding the residual plastic potential is important if we aim to re-open the critical period after insult. Particularly important is ocular dominance plasticity in amblyopia (Webber & Wood, 2005), a cortical deficit still without cure in adults, although recent advancements in training procedures are opening new hopes (Levi & Li, 2009; Sengpiel, 2014). Endorsing plasticity may increase the effectiveness of these treatments and preliminary data from our laboratory suggest that monocular deprivation of the amblyopic eye may indeed boost sensitivity of the deprived eye and improve its acuity (Lunghi et al., 2016) – exactly as observed in our BOLD results in normally sighted participants. Although sensory primary cortices are particularly resilient to plastic change in adult, our data demonstrate that two hours of abnormally unbalanced visual experience induce a functional reorganization of cortical circuits, particularly of the parvocellular pathway, leading to an alteration of basic visual perceptual abilities.

## Methods text

### EXPERIMENTAL MODEL AND SUBJECT DETAILS

#### Human subjects

Experimental procedures are in line with the declaration of Helsinki and were approved by the regional ethics committee [Comitato Etico Pediatrico Regionale—Azienda Ospedaliero-Universitaria Meyer—Firenze (FI)] and by the Italian Ministry of Health, under the protocol “Plasticità e multimodalità delle prime aree visive: studio in risonanza magnetica a campo ultra alto (7T)”.

Twenty healthy volunteers with normal or corrected-to-normal visual acuity were examined (8 females and 12 males, mean age = 27 years) after giving written informed consent.

### METHOD DETAILS

#### Experimental design

Each participant underwent two scanning sessions separated by two hours, during which they were subject to the short-term monocular deprivation procedure described below. Just before each scanning section, their binocular rivalry was measured psychophysically. One (male) participant was excluded because of strong eye dominance tested with binocular rivalry before the deprivation. This left 19 participants (8 females and 11 males) whose complete datasets were entered all analyses. Sample size was set to enable testing for correlations between neuroimaging and psychophysical data. Previous work (Lunghi, Emir, et al., 2015) reveals a correlation between MR spectroscopy data and binocular rivalry measures r = 0.62 (or higher), which implies a minimum of 17 participants to detect a significant correlation at 0.05 significance level, with test power of 80%(Lachin, 1981).

#### Short-term Monocular Deprivation

Monocular deprivation was achieved by patching the dominant eye for 2 hours. The operational definition of dominant eye applied to the eye showing the longer phase durations during the baseline binocular rivalry measurements. Like in previous studies (Binda & Lunghi, 2017; Lunghi et al., 2011; Lunghi et al., 2013), we used a translucent eye-patch made of plastic material allowing light to reach the retina (attenuation 0.07 logUnits, at least 3 times smaller than the threshold for discriminating a full-field luminance decrement (Knau, 2000) and more than ten times smaller than the minimum photopic luminance decrement required for shifting the spatial (Van Nes & Bouman, 1967) or temporal contrast sensitivity function (Kelly, 1961)). The patch prevents pattern vision, as assessed by the Fourier transform of a natural world image seen through the eye-patch. During the 2 hours of monocular deprivation, observers were free to read and use a computer.

#### Binocular Rivarly

Binocular rivalry was measured in two short sessions (each comprising two runs of 3 minutes each), immediately before the Pre- and Post-deprivation MR sessions, in a quiet space adjacent to the MR control room. Visual stimuli were created in MATLAB running on a laptop (Dell) using PsychToolbox (Brainard, 1997), and displayed on a 15-inch monitor (BenQ). Like in (Lunghi, Emir, et al., 2015), observers viewed the visual stimuli presented on the monitor at a distance of 57 cm through anaglyph red-blue goggles (right lens blue, left lens red). Responses were recorded with the computer keyboard by continuous alternate keypresses. Visual stimuli were two oblique orthogonal red and blue gratings (orientation: ±45°, size: 3°, spatial frequency: 2 cpd, contrast 50%), surrounded by a white smoothed circle, presented on a black uniform background in central vision. Peak luminance of the red grating was reduced to match the peak luminance of the blue one using photometric measures. All included participants had typical binocular rivalry dynamics, with low percentage of mixed percepts (reported for 8.5 ± 2.04% of time on average). Only one participant experienced of mixed percepts for more than 20% of time (exactly for 31.2%) and his data are in line with the others’.

#### Stimuli for fMRI

Visual stimuli were projected with an MR-compatible goggle set (VisuaStimDigital, Resonance Technologies, Los Angeles, USA), connected to a computer placed in the control room. The goggles covered a visual field of approximately 32 × 24 deg, with a resolution of 800 × 600 pixels, mean luminance 25 cd/m^2^; the images in the two eyes were controlled independently.

During all functional MRI scans participants were instructed to maintain central fixation on a red point (0.5 degrees) that was constantly visible at the center of the screen. Bandpass noise stimuli were white noise images filtered to match the spatial frequency tuning of neurons in the visual cortex (Morrone & Burr, 1988). We generated a large white noise matrix (8000 × 6000) and filtered it with a two-dimensional circular bandpass filter *Bp* defined by Eq. 1:

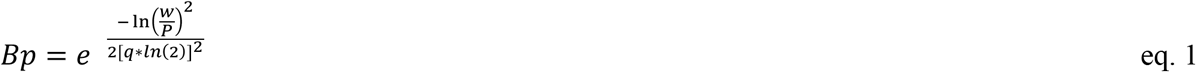

where *P* is the peak spatial frequency, *q* is the filter half-width at half maximum in octaves. We generated five band-pass noise stimuli, by setting *q* = 1.25 octaves and *P* = 0.1 cpd, 0.2 cpd, 0.4 cpd, 1.1 cpd, 2.7 cpd. Each stimulus was presented for a block of 3TRs, during which the image was refreshed at 8Hz (randomly resampling a 800 × 600 window from the original matrix). Stimuli were scaled to exploit the luminance range of the display, and this yielded very similar RMS contrast values (shown in supplementary Fig. S2). Stimulus blocks were separated by 4TRs blanks, consisting of a mid-level gray screen. The five band-pass noise stimuli blocks were presented in pseudo-random order, twice per run, for a total of 70 TRs. In each run, stimuli were only presented to one eye, while the other was shown a mid-level gray screen. Each eye was tested once, before and after deprivation.

Immediately upon application of the monocular patch, we performed two additional scans to perform retinotopic mapping of visual areas. Meridian and ring stimuli were presented monocularly (to the non-patched eye) and were defined as apertures of a mid-level gray mask that uncovered a checkerboard pattern, 1 deg at 1 deg eccentricity to 2.5 deg at 9 deg eccentricity, rotating and contracting at a rate of one check per second. Meridians were defined by two 45° wedges centered around 0° or around 90°. The horizontal and vertical meridian were presented interchangeably for 5 TRs each (without blanks) and the sequence was repeated 6 times for a total of 60 TRs. Rings partitioned screen space into six contiguous eccentricity bands (0-0.9 deg, 0.9-1.8 deg, 1.8-3.3 deg, 3.3-4.7 deg, 4.7-6.48 deg, 6.48-9 deg). Odd and even rings were presented in two separate runs. In each run, the three selected rings and one blank were presented in random order for 5 TRs each, and the sequence was repeated (with different order) 6 times for a total of 120 TRs.

#### MR system and sequences

Scanning was performed on a Discovery MR950 7 T whole body MRI system (GE Healthcare, Milwaukee, WI, USA) equipped with a 2-channel transmit driven in quadrature mode, a 32-channel receive coil (Nova Medical, Wilmington, MA, USA) and a high-performance gradient system (50 mT/m maximum amplitude and 200 mT/m/ms slew rate).

Anatomical images were acquired at 1 mm isotropic resolution using a T1-weighted magnetization-prepared fast Fast Spoiled Gradient Echo (FSPGR) with the following parameters: TR = 6 ms, TE = 2.2 ms. FA=12 deg, rBW = 50kHz, TI = 450 ms, ASSET = 2.

Functional images were acquired with spatial resolution 1.5 mm and slice thickness 1.4 mm with slice spacing = 0.1 mm, TR = 3 ms, TE = 23ms, rBW = 250 kHz, ASSET = 2, phase encoding direction AP-PA. No resampling was performed during the reconstruction. For each EPI sequence, we acquired 2 additional volumes with the reversed phase encoding direction.

### QUANTIFICATION AND STATISTICAL ANALYSIS

#### ROI definition

Areas V1, V2 and V3 were manually outlined for all participants using retinotopic data projected on surface models of white matter. The V1/V2 boundary was traced from the vertical/horizontal meridian flip superior/inferior to the calcarine sulcus, and the V2/V3 border and V3 border from the subsequent opposite flips. Areas V4, V3a and MT were defined based on the cortical parcellation atlas by Glasser et al. (Glasser et al., 2016). V1, V2, V3, V4 and V3a ROIs were further restricted to select the representation of our screen space. Specifically, the anterior boundaries were defined based on activation from most peripheral (6.48°-9°) ring stimuli of the retinotopic mapping scans; in addition, vertices were only included in the analysis if their preferred eccentricity (estimated through Population Receptive Field modelling, see below) was larger than 1, since no reliable mapping could be obtained for the central-most part of the visual field.

#### Pre-processing of imaging data

Analyses were performed mainly with Freesurfer v6.0.0, with some contributions of the SPM12 and BrainVoyager 20.6 and FSL version 5.0.10 (Jenkinson, Beckmann, Behrens, Woolrich, & Smith, 2012) packages.

Anatomical images were corrected for intensity bias using SPM12 (Friston, 2007) and processed by a standard procedure for segmentation implemented in Freesurfer (recon-all: Fischl et al., 2002). In addition, each hemisphere was aligned to a left/right symmetric template hemisphere (fsaverage_sym: Greve et al., 2013).

Functional images were corrected for subject movements (Goebel, Esposito, & Formisano, 2006) and undistorted using EPI images with reversed phase encoding direction (Brain Voyager COPE plug-in Jezzard & Balaban, 1995). We then exported the preprocessed images from BrainVoyager to NiFTi format. These were aligned to each participant’s anatomical image using a boundary based registration algorithm (Freesurfer *bbergister* function) and projected to the cortical surface of each hemisphere. All analyses were conducted on data in the individual subject space. In addition, for visualization purposes, we also aligned the results of timecourse analyses (GLM and subsequent pRF and spatial frequency tuning estimates) to the left/right symmetric template hemisphere. Averaged results across the 18×2 hemispheres are shown in the maps of Fig. 1B, Fig. 5A and Supplementary Fig. S1.

#### GLM analysis of fMRI data

General Linear Model analysis was performed with in-house MATLAB software (Mathworks, version R2016b). We assumed that fMRI timecourses result from the linear combination of N predictors: boxcar functions representing stimulus presence/absence (one per stimulus type) convolved by a standard hemodynamic response function (see Eq. 2), plus two nuisance variables (a linear trend and a constant). We modeled the hemodynamic response function as a gamma function h(t):

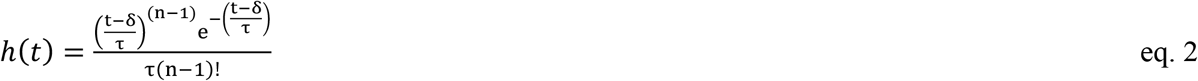

with parameters *n*=3, *t*=1.5 s, and d=2.25 s (Boynton, Engel, Glover, & Heeger, 1996). Beta weights of the stimuli predictors were taken as estimates of the BOLD response amplitude and normalized by the predictor amplitude to obtain a measure that directly corresponds to % signal change; beta weights were also scaled by an error measure to obtain t-values, following the same procedure as in (Friston et al., 1994). Computing BOLD responses for each individual vertex of the cortical surface leads to up-sampling the functional data (each 1.5 × 1.5 × 1.5 mm functional voxel projecting on an average of 3 vertices). We ensured that this does not affect our statistical analyses by first averaging data from all vertices within a region of interest (e.g. V1), thereby entering all ANOVAs with a single value per subject and region of interest.

#### Population Receptive Field mapping

The pRFs of the selected voxels were estimated with custom software in Matlab, implementing a method related to that described by Dumoulin and Wandell (Dumoulin & Wandell, 2008). We modeled the pRF with a 1D Gaussian function defined over eccentricity, with parameters *ε* and *σ* as mean and standard deviation respectively, and representing the aggregate receptive field of all neurons imaged within the vertex area. We defined the stimulus as a binary matrix S representing the presence of visual stimulation over space (here, eccentricity between 0 and 10 deg with 40 steps per deg) for each of 6 ring stimuli. We used the results of our GLM analysis to estimate the vertex response to each of our 6 rings (as t-values; using beta values yields very similar results). We assumed that each vertex response is the linear sum over space (eccentricity) of the overlap between the pRF of the voxel and the input stimulus, which is mathematically equivalent to the matrix multiplication between the stimulus and the pRF.

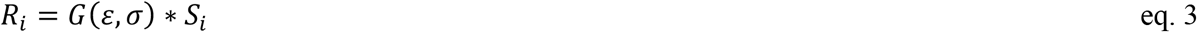

where *i* is the index to ring number and varies between 1 and 6.

We used this equation to predict the response to our six rings for a large set of initial pRF parameters; for each vertex, we measured the correlation (our goodness-of-fit index) between the predicted response and the observed t-values. If the highest correlation was < .7 the vertex was discarded; otherwise, the parameters yielding the highest correlation were used to initialize a nonlinear search procedure (MATLAB simplex algorithm), which manipulated *ε* and *σ* to maximize goodness-of-fit, with the constraint that *ε* could not exceed 20 deg or be smaller than 1 deg, and *σ* could not be smaller than .1 deg. Successful fits were obtained for 72.00 ± 1.86% of V1 vertices, for which the initial coarse grid search gave a correlation > 0.7 and the nonlinear search settled within the constraints. All analyses (on average and distribution of responses and tuning parameters) considered the sub-region of V1 for which a successful fit was obtained. We used *ε* to estimate the preferred eccentricity of each vertex.

The main modifications of our procedure relative to that described by Dumoulin and Wandell (Dumoulin & Wandell, 2008) are the following: (a) fMRI data were acquired in a block design with only six stimulus types (six eccentricity bands) rather than varying stimulus position at each TR; this allowed us to use a standard GLM approach to estimate each vertex response to the six stimuli (assuming a standard hemodynamic response function) and then use the pRF model to predict these six time-points – much faster than predicting the full fMRI series of 120×2 TRs; (b) our stimuli and consequently our pRFs were defined in one dimension (eccentricity) – whereas the standard pRF is defined in 2D, eccentricity and polar angle (or Cartesian x and y); (c) we maximized the correlation between the predicted and observed fMRI response time-courses rather than minimizing the root mean square error; this eliminates the need to estimate a scale factor to account for the unknown units of the BOLD signal.

#### Population Tuning for Spatial Frequency

Using a similar logic, we also estimated the population tuning for Spatial Frequency, which represents the aggregate Spatial Frequency tuning of the population of neurons imaged within each vertex area. We modeled the population tuning using a family of functions that includes the psychophysical Contrast Sensitivity Function (CSF) and can be specified by the following one-parameter equation (Difference-of-Gaussians):

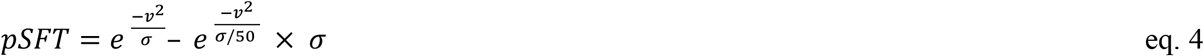

Like we did for the pRF mapping, we defined a stimulus matrix S representing the Fourier spectra of our five bandpass noise stimuli, i.e. the energy of visual stimulation in the frequency domain (here, between 0.03 cpd and 12.5 cpd) for each stimulus. We used the results of our GLM analysis to estimate the vertex response to each of our five bandpass noise stimuli (as t-values; using beta values yields very similar results). We assumed that each vertex response is the linear sum over frequency of the overlap between the pSFT of the voxel and the input stimulus, which is mathematically equivalent to the matrix multiplication between the stimulus and the pSFT.

Like for pRFs, we estimated the best-fit *σ* parameter of each vertex pSFT with a two-step procedure: a coarse-grid search followed by the simplex search. We used the matrix multiplication of the pSFT and the stimulus to predict the response to our five bandpass noise stimuli for a large set of initial *σ* values (between 1 and 1,000 in 100 logarithmic steps); for each vertex, we measured the correlation (our goodness-of-fit index) between the predicted response and the observed t-values. If the highest correlation was < .5, the voxel was discarded, otherwise the parameter yielding the highest correlation were used to initialize a nonlinear search procedure (MATLAB simplex algorithm), which manipulated *σ* to maximize goodness-of-fit, with the constraint that *σ* could not be smaller than .3 and larger than 10,000. Successful fits were obtained for 88.84 ± 1.28% of V1 vertices for which we obtained a successful eccentricity fit (86.77 ± 1.25% of all V1 vertices).

We express the *σ* parameter in terms of the high-spatial frequency cutoff of the filter (highest spatial frequency at half maximum), *SFco* for each vertex:

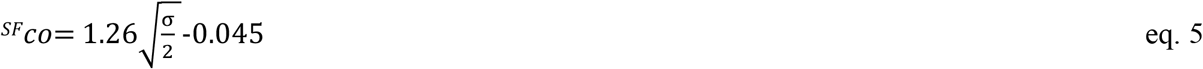

#### Indices defining the effect of deprivation

We computed the effects of short-term monocular deprivation on both the dynamics of binocular rivalry and our fMRI results, estimating the degree to which the two measures are correlated. In all cases, the same equation was applied to psychophysical and fMRI data.

The first index, called “Deprivation Index” or DI_psycho_ and DI_BOLD_ is given by eq. 6

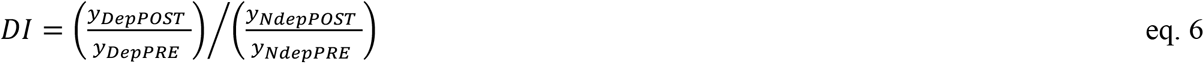

For psychophysics, y = mean duration of Binocular Rivalry phases of the Dep or Ndep eye, during the PRE-or POST deprivation sessions; for fMRI, y = mean BOLD response across V1 vertices to stimuli in the Dep or Ndep eye, during the PRE-or POST-deprivation sessions.

The second index, called “Deprived-eye change” or DepC_psycho_ and DepC_cutoff_ is given by eq. 7

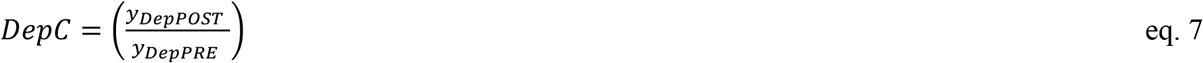

For psychophysics, y = mean duration of Binocular Rivalry phases of the Dep eye, during the PRE-or POST deprivation sessions. For fMRI, y = mean spatial frequency cut-off across V1 vertices estimated for stimuli in the Dep eye, during the PRE-or POST-deprivation sessions.

#### Statistics

Data from individual participants (mean binocular rivalry phase durations or mean BOLD responses/pRF/pST across V1 or V2 vertices) were analyzed with a repeated measure ANOVA approach, after checking that distributions do not systematically deviate from normality by means of the Jarque-Bera test for composite normality (Matlab *jbtest* function, p-values given in the relevant figures). F statistics are reported with associated degrees of freedom and p-values in the Results section, in the form: F(df,df_err_) = value; p = value. Post-hoc paired t-tests comparing conditions follow the ANOVA results, in the form: t(df) = value, p = value. Associations between variables are assessed with Pearson product-moment correlation coefficient, reported in the form: r(n) = value, p = value. Aggregate subject data (i.e. vertices pooled across participants and hemispheres) were typically non-normally distributed and thereby were analysed with non-parametric tests. The Wilcoxon sign-rank test was used for comparing medians, and results are reported in the form: z = value, p = value.

## DATA AND SOFTWARE AVAILABILITY

Data and software are available on Dryad (linked to the current eLife submission).

## Acknowledgments

This research was supported by the European Research Council under the European Union’s Seventh Framework Programme (FPT/2007-2013) under grant agreement number 338866 (P.B., C.L., and M.C.M.) and under ERA-NET project “Neuro-DREAM” (C.L., and M.C.M.) and by the European Union’s Horizon 2020 Research and Innovation Programme under the Marie Sklodowska-Curie grant agreement number 641805 (J.W.K.) and by the Italian Ministry of University and Research under the project PRIN2015 (M.C.M) and by Fondazione Roma under the Grants for Biomedical Research: Retinitis Pigmentosa (RP)-Call for proposals 2013-“Cortical Plasticity in Retinitis Pigmentosa: an Integrated Study from Animal Models to Humans”. The authors would like to thank Mauro Costagli for help with the data acquisition.

## Author contributions

P.B., C.L., and M.C.M. designed the experiments. J.W.K., P.B., C.L. and L.B. performed the experiments and M.T. supervised the 7T scanning. P.B. and J.W.K. analyzed the results. P.B. and M.C.M. wrote the manuscript. All authors revised the manuscript.

## Materials and correspondence

Correspondence should be addressed to Maria Concetta Morrone

